# Relative energetic economy of cleistogamous selfing in three populations of the perennial *Ruellia humilis*

**DOI:** 10.1101/2023.05.23.541962

**Authors:** Tatyana Y. Soto, Nicholas A. Ryan, Christopher G. Oakley

## Abstract

**Premise of research:** What maintains mixed selfing and outcrossing is an enduring mystery in evolutionary biology. Cleistogamy, where individuals produce both potentially outcrossing chasmogamous and obligately selfing cleistogamous flowers, provides an ideal framework in which to study the evolutionary forces maintaining mixed-mating. Despite their promise for providing insight into mating system evolution, few studies of cleistogamous species have comprehensively investigated the relative costs and benefits of cleistogamous selfing.

**Methodology:** We quantified the necessary components to calculate the relative energetic cost of reproduction of the different flower types in three natural populations of the perennial *Ruellia humilis* Nutt (Acanthaceae). These components include flower dry mass, fertility (fruit set), and seed mass and number per fruit. We also measured pollen-ovule ratios for both flower types as another measure of relative energetic investment. We additionally tracked phenology of the two flower types and used the proportion of chasmogamous flowers to estimate maximum potential outcrossing rates.

**Pivotal Results:** We found that the energetic cost of reproduction via cleistogamous flowers was about 4-10 times less than that of reproduction via chasmogamous flowers, and this energetic economy was from both reduced mass and increased fertility of cleistogamous flowers. Pollen ovule ratios in cleistogamous flowers were one third to one half those of chasmogamous flowers, providing additional support for their energetic economy. Maximum potential outcrossing rates in these populations based on chasmogamous flower production were between 43-61%, but chasmogamous flowers can autogamously self at rates of 33-75%.

**Conclusions:** The relative energetic economy of cleistogamous flowers suggest that progeny from chasmogamous flowers would have to have 4-10 times greater relative fitness to explain their evolutionary maintenance. These values are likely even greater considering the reduced investment in pollen production in cleistogamous flowers. Ongoing work will quantify potential advantages of chasmogamous flowers due to inbreeding depression and heterosis.

## Introduction

The persistence of mixed mating, reproduction by a combination of selfing and outcrossing, is an old and unresolved mystery in plant evolutionary biology. The evolution of mating systems depends on the balance between the costs and benefits of self-fertilization. Inbreeding depression, or the reduction in fitness of inbred relative to outbred progeny, plays a central role as a cost of increased selfing (Lande and Schemske, 1985; Charlesworth and Charlesworth, 1987; Goodwillie et al., 2005; Winn et al., 2011). However, self-fertilization may be beneficial in providing reproductive assurance with limited pollinator or mate availability (Baker, 1955; Lloyd, 1992; Kalisz et al., 2004). Additionally, reduced flower size in selfing species can provide a benefit to selfing via greater energetic economy (Randle et al., 2009; Grossenbacher and Whittall, 2011). One additional consideration in understanding the adaptive significance of mixed mating is that some self-fertilization may simply be accidental. For example, large floral display sizes may attract more pollinators but also lead to greater selfing among different flowers on the same plant (de Jong et al., 1993; Harder and Barrett, 1995). An ideal system for investigating the benefits of selfing would be one with autonomous self-fertilization and the ability to quantify the relative energetic economy of selfing compared to outcrossing. The floral heteromorphism in cleistogamous species provide such an opportunity, yet these species have received less attention in the plant mating system literature (Goodwillie et al., 2005; Oakley et al., 2007; Winn et al., 2011).

Individual plants of cleistogamous species produce both potentially outcrossing chasmogamous (CH) flowers and cleistogamous (CL) flowers that are closed and obligately selfing. Following historical precedent (Lord, 1981), we refer to cleistogamy or cleistogamous to refer to the syndrome of producing both flower types, and cleistogamous (CL) flowers to refer to the selfing flower type. Mixed production of CH and CL flowers occurs in over fifty families, and has evolved independently multiple times from an ancestral state of only CH flowers, with only three documented evolutionary losses of CH flowers (Culley and Klooster, 2007). The repeated independent origins, broad taxonomic occurrence, and persistence of both obligately selfing and potentially outcrossing flowers on the same individual plant provides strong evidence that cleistogamy is an adaptive strategy for mixed mating (Culley and Klooster, 2007; Oakley et al., 2007).

The production of both flower types suggesting adaptive mixed mating is however at odds with results from studies on the costs and benefits of CL selfing. Taken together this body of work indicates that it is difficult to explain the maintenance of CH outcrossing (Oakley et al., 2007). Evidence for a cost of CL selfing suggests that it is weak. Relative components of fitness of progeny derived from CL selfing is similar on average (across about 20 studies in 14 species) to that of progeny derived from CH flowers (reviewed in Oakley et al., 2007), and estimates of inbreeding depression in cleistogamous species fall below the 50% threshold expected to prevent the evolution of complete selfing (Culley, 2000; Oakley et al., 2007; Oakley and Winn, 2008; Ansaldi et al., 2019). One benefit of CL flowers is that they have higher rates of fruit set, or fertility, compared to CH flowers, and may therefore assure reproduction in the absence of pollinators (Culley, 2002; reviewed in Oakley et al., 2007; Ansaldi et al., 2019). Additionally, CL flowers may be more energetically favorable than CH flowers because they are highly reduced in size (Schemske, 1978; Waller, 1979; Oakley et al., 2007) and typically lack pigmentation and nectar (Lord, 1981). However, our current knowledge on the relative benefit of CL flowers is mostly pieced together from estimates of isolated components of costs and benefits across single populations of many different species. Measuring a complete suite of traits related to the energetic cost of reproduction of each flower type in multiple population of the same system will permit us to better estimate how strong selection for outcrossing must be to maintain the production of CH flowers in spite of the benefits that CL flowers provide (Oakley et al., 2007).

To quantify relative energetic costs of reproduction of the two flower types, estimates of flower mass, fertility (fruit set), seed number per fruit, and seed weight for both CH and CL flowers are needed. Dry flower mass has been used as a proxy for resource investment to reproduction, and CL flowers can be 1 to 100-fold lighter than CH flowers (Schemske, 1978; Waller, 1979; Schoen, 1984; Sun, 1999; Oakley et al., 2007; Seguí et al., 2021). Fertility of CL flowers is typically high, whereas fertility of open pollinated CH flowers can range from 25% to 96% depending on the species (Oakley et al., 2007; Abdala-Roberts et al., 2014; Goodwillie et al., 2018). Higher fertility of CL flowers decreases the cost of reproduction because less energy is invested into flowers that may not result in seed set. Seed mass is directly related to the energetic cost of reproduction, and sometimes there are dramatic differences in seed mass produced by CH and CL fruits (Sun, 1999; Berg and Redbo-Torstensson, 2000; Albert et al., 2011; Cheplick, 2023). Finally, standardizing the cost of reproduction of each flower type by seed number per fruit is important to directly compare the two flower types (Oakley et al. 2007) if different flower type have different numbers of seeds (Culley, 2000; Albert et al., 2011; Ansaldi et al., 2019).

Another way to quantify relative energetic investment in reproduction of the two flower types is the ratio of pollen to ovules in a flower. Pollen is lightweight and may thus not contribute to detectable differences in flower mass, but producing male gametophytes is expected to be costly (Schemske, 1978; Schoen and Lloyd, 1984). Lower pollen:ovule is expected in cleistogamous flowers because of the efficiency of self-fertilization in the closed flowers, adding to their energetic benefit (Cruden, 1977). This is one explanation for why cleistogamy is common in grasses, where the costly investment in production of male gametophytes required for wind-pollination could be reduced via CL flowers (Campbell et al., 1983). The only two studies that we are aware of that have measured pollen-ovule ratios in both flower types of cleistogamous species have documented up to a 5-fold increase in pollen:ovule of CH flowers relative to CL flowers (Lord, 1980; Sun, 1999).

Relative phenology and degree of chasmogamy (proportion of all flowers that are CH) are two additional traits that can provide insight into the selective forces that may be involved in maintaining CH flowers. For example, one model (Schoen and Lloyd, 1984) frames the floral heteromorphism as a type of adaptive plasticity, where the plant matches the flower type to seasonal variation in order to maximize fitness. Thus, a pattern of producing different flower types in distinct seasons would then be a necessary, but not sufficient, condition for this hypothesis (Winn and Moriuchi, 2009; Stojanova et al., 2016). Information on the degree of chasmogamy likewise provides an upper limit on the population level outcrossing rate in the population, though the realized outcrossing rate may be much lower depending of rates self-fertilization in CH flowers (Oakley et al., 2007).

There are estimates of the individual traits discussed above in cleistogamous species (reviewed in Oakley et al., 2007), but few studies measure all of the necessary traits to quantify the relative energetic cost of reproduction of the two flower types (Schemske, 1978; Waller, 1979; Winn and Moriuchi, 2009) in the same system. Additionally, three of the four most complete estimates have only been done in a single population per species. Estimates from replicate populations ensures a more general characterization of the relative benefits of CL selfing. Here we quantify the necessary components for cost of reproduction: flower dry mass, fertility, seed weight, and seed number from three natural populations of *Ruellia humilis* Nutt (Acanthaceae). In addition to these components, we measured pollen-ovule ratios of both flower types as another proxy to energetic investment into reproduction. Lastly, we tracked the phenology of CH and CL production (distinct seasons vs. overlapping) and calculated the degree of chasmogamy in all three populations.

## Materials and Methods

### Study system

*Ruellia humilis* Nutt (Acanthaceae) is a short-lived perennial native to rich sandy soils of prairies in the midwestern United States. Like many species in this genus (Tripp, 2007), *R. humilis* is cleistogamous and individual plants produce both CH and CL flowers. This species is primarily pollinated by hawkmoths, but a decline in visitation to *R. humilis* has been documented (Heywood et al., 2017; Heywood et al., 2022). The anthers of both CH and CL flowers dehisce around dusk and flowers of both types last for less than a day. Selfing of CH flowers has been observed as the anthers brush against the stigma when the corolla tube falls off of the plant in the afternoon (Long and Uttal, 1962).

The number of native populations of *R. humilis* are limited because of widespread conversion of prairie habitat for agricultural land use. Remnant populations do occasionally persist along roadsides and cemeteries established on sandy soils, which serve as habitat for this and other prairie species. We identified three native populations in Indiana and Illinois from which we collected the source material for this work. Two cemetery prairie remnant populations Sand Ridge (40.4085° N, 86.9665° W) and St Mary’s (40.3952° N, 86.9172° W), and a roadside population Wilmington (41.3124° N, 88.1321° W).

### Germination of seeds for floral trait measurements

We collected seed from fruits of 20-23 maternal lines from these natural populations (unknown flower type). Seeds were germinated and plants were grown for a generation in the greenhouse to minimize maternal environment effects. The next generation of seeds were collected from 8-15 (mean = 10.33) maternal lines per population from CL fruits that developed in the greenhouse.

To grow plants for trait measurements, seeds were cold stratified on moist paper towels for 8 weeks in the dark at 4°C to break dormancy. After stratification, seeds were individually sown across six 285-cell flat inserts (volume = 7 cm^3^, Greenhouse Megastore, Danville, IL) filled with a 2:1 mixture of Berger BM2 potting soil (Berger Horticulture Products, Sulphur Springs, TX) and sand. After sowing, flats were placed in an incubator (Percival Scientific, Perry, IA) set at constant 28°C with 13-hour days with the maximum light intensity for this incubator, 65 μmoles/m^2^/sec. Once seedlings had two sets of true leaves, we transplanted them into 9 cm pots (volume = 245 cm^3^, Greenhouse Megastore Danville, IL) filled with the same sand/soil mixture and placed into a heated greenhouse (minimum temperature: 20°C). Four weeks after the initial transplant, we transplanted the plants again into larger pots (10.79cm x 10.79cm x 12.38cm, Greenhouse Megastore, Danville, IL) filled with the same sand/soil mixture and kept them in the greenhouse for the remainder of the experiment. Individuals began flowering nine weeks after sowing in April 2022.

### Traits for calculating the energetic costs of reproduction

For dry flower mass, we sampled 1-3 flowers of each type (CH and CL) on replicate plants from 8 to 11 maternal lines. Entire flowers (minus two anthers used for pollen-ovule ratios, see below) were harvested, placed in a coin envelope, and set in a drying oven at 60°C for ten days. The flowers were then weighed to the nearest 0.01 mg (model: ML104, Mettler Toledo, Columbus, OH). We estimated mean fertility of CL and CH flowers as autogamous fruit set by marking buds with uniquely colored thread for each individual plant. Autogamous fruit set in a pollinator free environment is an estimate of minimum fertility because outcrossing the field may increase CH fertility. After two weeks, fruit set was visually scored as 1 for flowers that produced fruit or 0 for those that did not. These fruits were collected in Fall 2022 once they had turned brown, and placed in coin envelopes and allowed to dehisce prior to weighing and counting seed number per fruit. Estimates of mean seed weight per fruit were then obtained by dividing total seed mass per fruit by the number of seeds per fruit.

To calculate the relative cost of reproduction by CL flowers compared to CH flowers (Oakley et al., 2007), we first averaged over CH and CL flowers for each replicate plant for dry mass, fertility, seed number per fruit and seed weight. We then averaged over all replicate individuals for each trait and flower type to obtain population level means (Table S1). Using these population averages, we calculated a population level cost of reproduction of each flower type using the following equation:

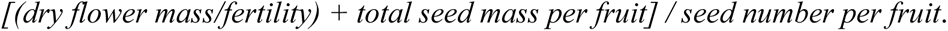

Fertility in the denominator of the first term accounts for the investment in flowers that do not yield seeds, and seed number per fruit standardizes the average cost of reproduction per fruit (in brackets) if CH and CL fruits produce different numbers of seeds. To get a relative cost of reproduction of the two flower types (CL/CH), we divided the cost of reproduction of CL flowers by that of CH flowers.

### Pollen ovule ratio

While the production of pollen and ovules does not contribute greatly to total flower mass, the production of these structures is expected to be costly. We therefore also quantified the ratio of pollen number to ovule number (P:O) as an alternative approach to quantifying relative costs of reproduction. For each of the same flowers harvested for dry mass (see above), we counted the number of ovules by dissection under a stereomicroscope. The total number of pollen grains (see below) was then divided by the number of ovules to get P:O per flower.

For pollen counts, we collected 2 of the 4 anthers from a given flower bud approximately two hours before anther dehiscence. The 2 anthers were placed in a microcentrifuge tube at room temperature for 12 hours. Once anthers had dehisced, we added 500μl to each centrifuge tube consisting of 450μl of a glycerol-ethanol-H_2_O solution, plus 50μl of 1% aniline blue to aid in visualizing pollen grains. We vortexed the tubes for one minute to release pollen grains from the anther sacs. After vortexing, five replicate 20μL aliquots of the suspended solution were pipetted onto five different ruled microscope slides with an etched grid of 64-2mm squares (Flinn Scientific, Batavia, IL), vortexing the microcentrifuge tube for thirty seconds between pipetting each aliquot. We counted pollen grains on the entire grid of each slide at 10x magnification under a light microscope and averaged the five separate counts for each flower. This metric represents the average number of pollen grains in a 20μL sample from half of the total pollen grains per flower suspended in a total of 500μL. Total pollen grains per flower was then estimated by multiplying this average by 25 (20μL x 25 = 500μL) to obtain the total number of pollen grains in the 500μl solution, and then multiplied by 2 to get the total pollen production from all four anthers on a flower.

### Flowering phenology and degree of chasmogamy

We quantified the timing of production of each flower type in *R. humilis* by counting the number of each of the CH and CL flowers on all flowering individuals every day for 22 weeks starting in April 2022. Because daily flower production was sparse, we summed weekly CH and CL production for each population over all replicate plants. Average number of CH and CL flowers produced per individual plant per population per week was then calculated for each population by dividing weekly totals by the number of flowering individuals. We also quantified the proportion of total flowers produced that were CH (CH:[CH+CL]) per individual plant by summing the total number of CH and CL flowers produced over the twenty-two-week period. This ratio gives the maximum potential outcrossing rate per population, though realized outcrossing rates may be less due to CH selfing.

### Statistical Analysis

For each trait, we tested for the effects of maternal population, flower type, and their interaction using an analysis of variance (ANOVA). All terms were treated as fixed effects because of the small number of maternal populations. Individual plant means by flower type were used for all traits, including flower dry mass, fertility, seed number per fruit, average seed weight per fruit, and P:O. Although the residuals were not perfectly normally distributed with equal variance for all traits, we used normal error distributions for all traits because qualitatively similar results were obtained both for models with alternative error distribution, and for non-parametric models, indicating that our results are robust to moderate violations of the assumptions of ANOVA.

In an ANOVA for any given trait, a significant effect of flower type indicates a difference between CH and CL flowers averaged over the effect of population. A significant interaction between these two effects indicates that differences between flower type depend on the maternal population. A significant effect of maternal population would indicate genetically based differences among populations averaged over the effect of flower type, though this is not of central interest to our objectives. For CH/total we used a single factor ANOVA to test for an effect of maternal population because this trait is a ratio of both flower types. All analyses were performed in JMP (v16.1).

## Results

### Traits measured on individual flower types

There were highly significant effects of both maternal population and flower type for flower dry mass, fertility, and pollen:ovule, and a significant interaction between maternal population and flower type for flower dry mass (Table 1). Dry mass of CL flowers was consistent across the three populations (Fig. 1A, Table S1), so the significant interaction effect for this trait was driven by modest variation among populations in CH flower dry mass. Flower mass of CL flowers was only 27% that of CH flowers for the Sand Ridge population, and this proportional decrease was more pronounced for the St. Mary’s (22%) and Wilmington (20%) populations (Fig. 1A, Table S1). Over both flower types, St. Mary’s had significantly greater flower mass than the other two populations. Fertility of CL flowers in proportion to CH flowers overall was approximately 50% greater, and though the interaction was not significant, this proportional difference is smaller in Sand Ridge than in the other two populations, 19% compared to 94% and 83% for St. Mary’s and Wilmington respectively (Fig. 1B, Table S1). Sand Ridge had greater fertility than the other two populations when considering both flower types combined. Pollen:ovule of CL flowers was 42% that of CH flowers overall (Fig. 2, Table S1). Although there was no significant interaction for pollen:ovule, there was some variation among populations (range = 33-50%) in relative pollen:ovule of the two flower types (Fig. 2, Table S1). Over both flower types, Sand Ridge had significantly greater pollen:ovule than the other two populations.

**Figure 1.**
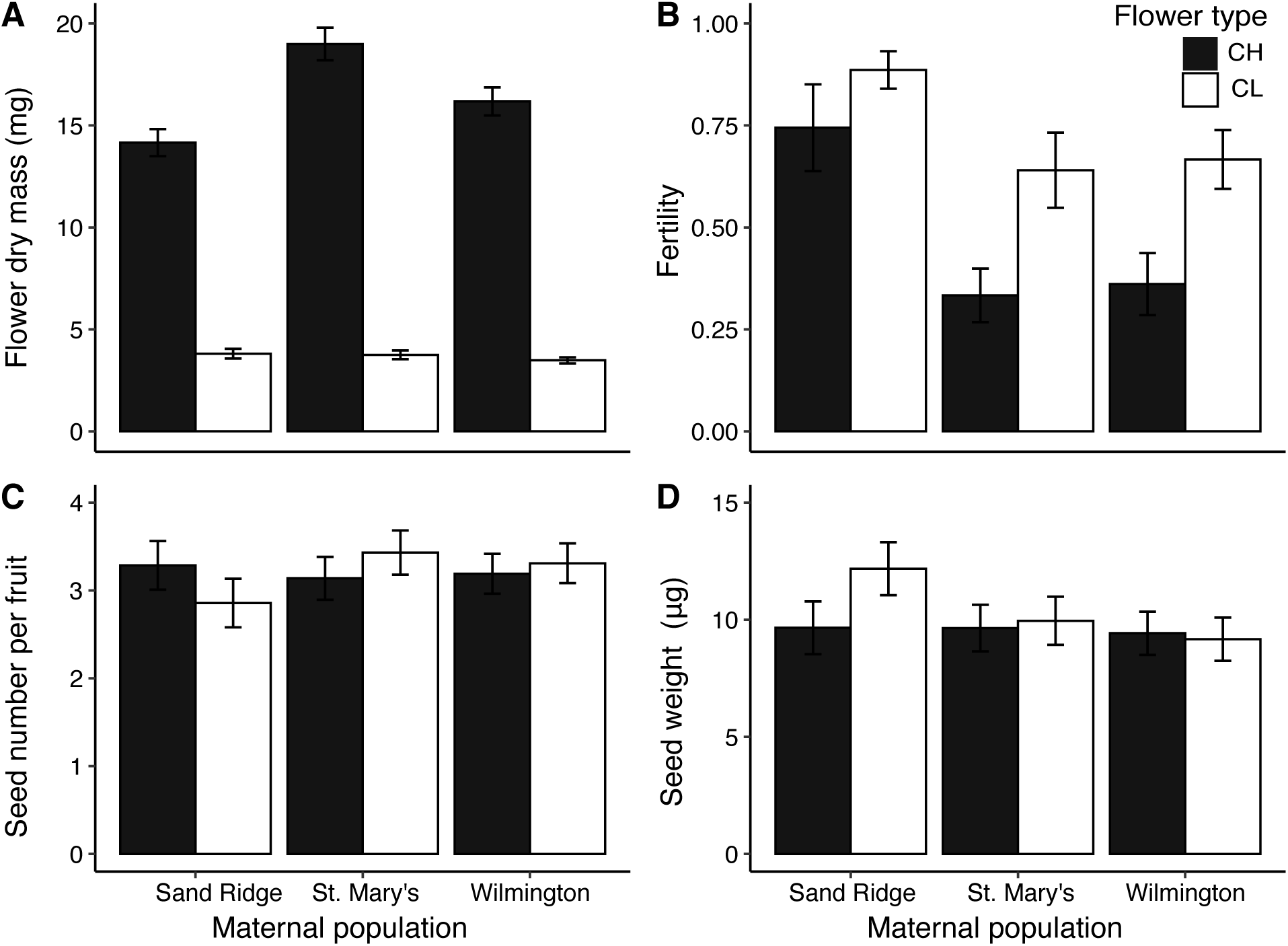
Mean trait values of chasmogamous (CH – black bars) and cleistogamous (CL – open bars) flowers by maternal population. **A.** Mean flower dry mass (mg). **B.** Mean fertility (probability of fruit set) averaged over multiple flowers of each type on a single individual plant. **C.** Mean seed number per fruit. **D.** Mean seed weight (μg). Error bars are +/-1 SE.

**Figure 2.**
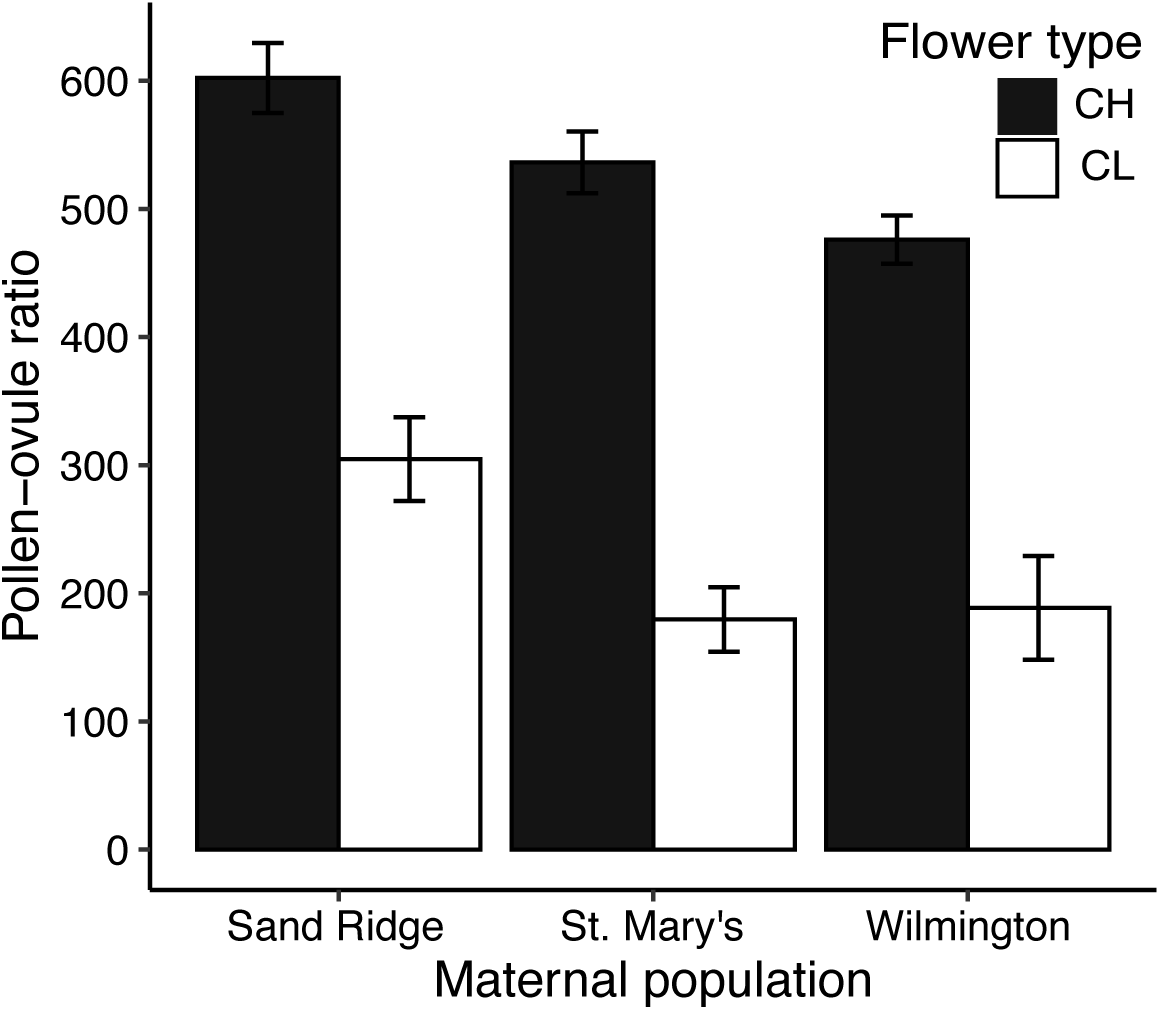
Mean pollen to ovule ratio of chasmogamous (CH – black bars) and cleistogamous (CL – open bars) flowers by maternal population. Error bars are +/-1 SE.

**Table 1.** ANOVA results for the effect of maternal population (3 populations), flower type (CH vs. CL), and their interaction on flower dry mass, fertility, mean seed number per fruit, mean seed weight per fruit, and pollen ovule ratio, and the ratio of CH flowers to the total number of flowers. F-ratios and denominator degrees of freedom are given for all traits.

| Trait | df | Population |  | Flower type |  | Population*type |  |
| --- | --- | --- | --- | --- | --- | --- | --- |
|  |  | F | P | F | P | F | P |
| Flower dry mass | 152 | 8.0 | <0.001 | 661.1 | <0.001 | 8.0 | <0.001 |
| Fertility | 126 | 9.9 | <0.001 | 15.6 | <0.001 | 0.7 | 0.511 |
| Seed number per fruit | 103 | 0.3 | 0.690 | 0.0 | 0.978 | 1.0 | 0.370 |
| Seed weight per fruit | 103 | 1.2 | 0.293 | 1.1 | 0.306 | 0.9 | 0.389 |
| Pollen:ovule | 116 | 9.5 | <0.001 | 171.1 | <0.001 | 0.9 | 0.424 |
| CH:total flowers | 121 | 8.1 | <0.001 | n/a | n/a | n/a | n/a |

Seed traits can be important components of the relative cost of reproduction by the two flower types if there are large differences in seed size and/or weight between CH and CL fruits. However, results for average seed number per fruit and average seed weight per fruit confirm initial observations that there were no differences with respect to flower type. In fact, there were no significant differences for these traits for any of the model terms (Fig. 1C&D, Table 1). Average seed number per fruit ranged between 2.8-3.4 across populations and cross types (Fig. 1 C, Table S1). Average seed weight per fruit was between 9.2-9.9μg across populations and cross types, with the exception of CL seeds in Sand Ridge which were somewhat larger (12.1 μg).

### Energetic costs of reproduction

Means for each flower type for flower mass, flower fertility, seed number per fruit, and the mass of those seeds are all components of the energetic costs of reproduction. Costs of reproduction via CL flowers was consistent across populations and ranged from 1.5-1.7 mg/seed (Table S1). Cost of reproduction via CH flowers was more variable among populations and ranged from 5.8 mg/seed in Sand Ridge, to 14.1 mg/seed in Wilmington, to 18.2 mg/seed in St. Mary’s (Table S1). Reduced overall cost of CH reproduction in Sand Ridge appears to be driven both by greater relative fertility (i.e., greater autogamous selfing) of CH flowers (Fig. 1B) and reduced relative mass of CH flowers (Fig. 1A). The relative (CL/CH) cost of reproduction in Sand ridge (0.26) was more than 2-fold higher than both St. Mary’s (0.09) and Wilmington (0.11). Because these derived variables are calculated at the population level, we were not able to perform a formal statistical analysis. However, results for individual components of cost of reproduction (Fig. 1) suggest that the difference between Sand Ridge and the other two populations is meaningful.

### Flowering phenology and degree of chasmogamy

Patterns of relative phenology of CH and CL flowers were qualitatively different between populations. In Sand Ridge, CH flower production started early and peaked by the third week of flowering, with CH flower production about 9-fold greater than CL flower production during the third week. This was followed by mixed production of both CH and CL flowers at a rate of about 2 flowers per individual per week of each flower type during weeks 4-14 (Fig. 3A). After week 14, CH flower production declined while CL production remained constant or increased slightly. In St. Mary’s, individuals started flowering two weeks later compared to the other populations (Fig. 3). In this population, CH flowers were almost exclusively produced for the first six weeks (Fig. 3B), peaking around week 7 where CH flower production was about 8-fold greater than CL flower production. Production of CL flowers in this population began increasing approximately eight weeks after first flowering, with mixed production of both flower types until week 16, and a shift towards almost exclusive CL production afterwards. In Wilmington, both CH and CL flower production began in the first week, and overall flower production was more consistently low across the 22 weeks compared to the other populations. Peak CH flower production occurred in week 1 and maximal CH production was about ½ that of the other two populations (Fig 3C). Production of both CH and CL flowers in a ratio about 3:1 continued until week 17, when CH production decreased, and CL production increased slightly. For the average ratio of CH flower number to total flower number over the entire flowering period, which represent the maximum potential outcrossing rates, we found values of 0.43 for Sand Ridge, 0.61 for St. Mary’s, and 0.52 for Wilmington. The value for Sand Ridge was significantly less than that of the other populations (Table 1, Fig. 4).

**Figure 3.**
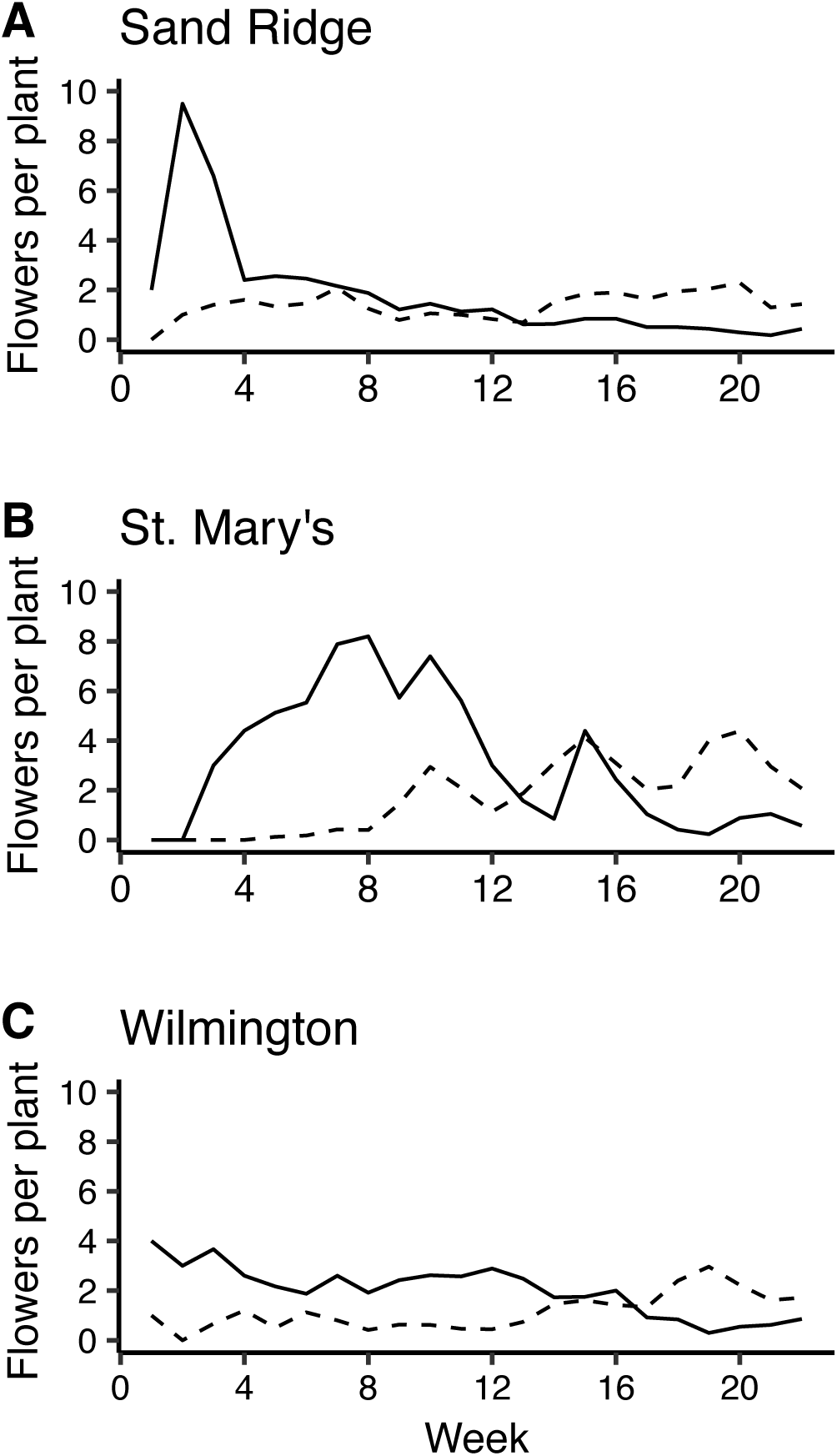
Average number of chasmogamous (CH – solid line) and cleistogamous (CL – dashed line) flowers produced (per individual per week) over a twenty-two-week period for the three maternal populations (A. Sand Ridge, B. St. Mary’s, C. Wilmington).

**Figure 4.**
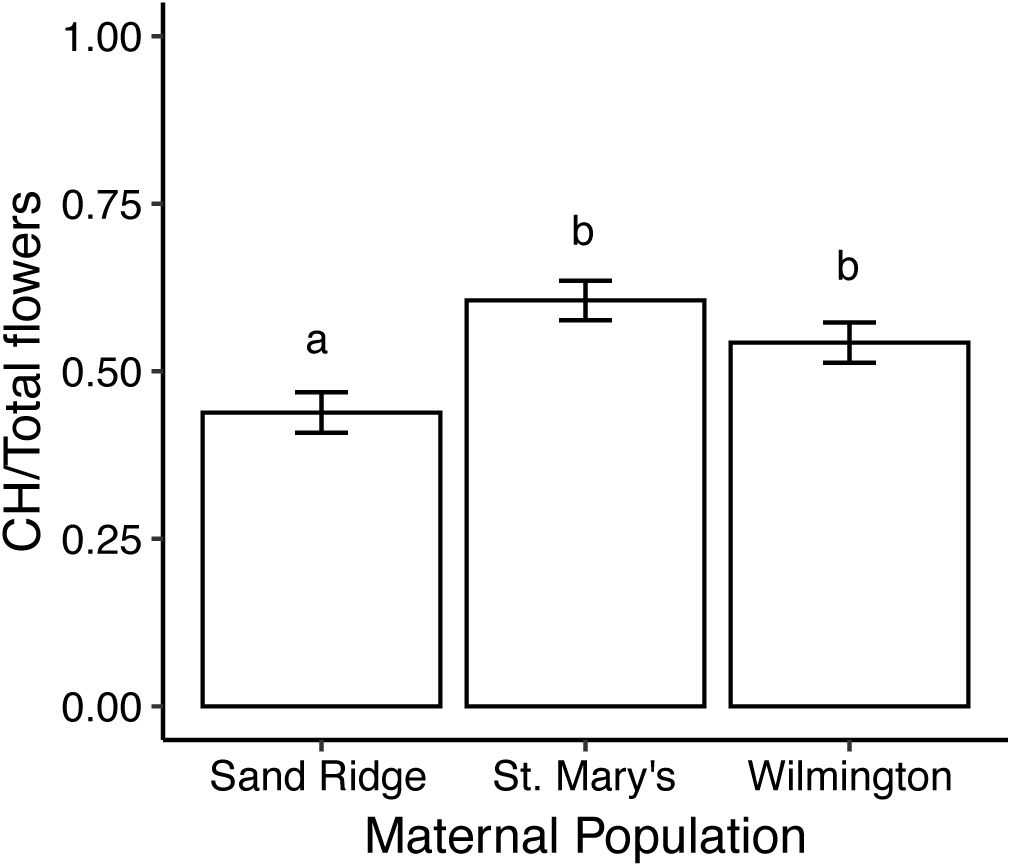
Mean proportion of total flower production that was chasmogamous by maternal populations. Error bars are +/-1 SE. Bars that do not share a letter are significantly different from one another.

## Discussion

The continued maintenance of CH flowers in cleistogamous species is a longstanding evolutionary enigma. However, only a few studies have collected all the information needed to understand the magnitude of selection required to maintain CH flowers, and only one study has done so for multiple populations. Overall, we found that the energetic cost of reproduction via CL flowers was between about 4-10 times less than that of CH reproduction, and this energetic economy comes both from reduced flower mass and increased fertility of CL flowers. Pollen ovule ratios in CL flowers were one third to one half those of CH flowers, providing additional support for the energetic economy of CL flowers. Finally, we found that about one half of the total flowers produced by these populations are CH, and these flowers are capable of autogamously selfing at rates of 33-75%.

### Traits for calculating the energetic costs of reproduction

We found that both flower dry mass and fertility were lower for CL compared to CH flowers. Dry weight of CL flowers relative to CH flowers ranged from 0.20-0.27 in *R. humilis*, which is somewhat lower than the average, but within the range, previously reported for cleistogamous species. Oakley et al. (2007) reviewed studies of relative energetic costs of the different flower types and reported estimates of relative flower mass from 6 species with a mean of 0.38 (range 0.01-0.9). A recent study in *Viola jaubertiana* reported a value of 0.29 (Seguí et al., 2021). For relative fertility of CL flowers, we found considerable variability between populations, with a range of 1.19-1.94. Values for relative fertility previously reported for 13 species ranged from 0.43-4.89 with a mean of 1.80 (Oakley et al., 2007) after excluding the extreme value of 19.2 for *Scutellaria indica* (Sun, 1999). A recent estimate of 14.3 for *V. jaubertiana* is likewise extreme (Seguí et al., 2021), but suggests that considerable variation in the relative biomass of CL flowers among species. As with relative flower mass, the values for relative fertility observed here are within the range of values previously observed, though the degree of interpopulation variation in *R. humilis* is noteworthy. In particular, the high absolute fertility of CH flowers in plants from Sand Ridge indicates a greater potential for autogamous selfing of CH flowers in this population. This result agrees with previous observations that CH flowers of *R. humilis* are capable of delayed selfing (Long and Uttal, 1962; Heywood et al., 2017). Although our estimates of CH fertility are conservative because outcrossing is not possible in the greenhouse. Our estimates of CH fertility in *R. humilis* are similar to field estimates of fertility of CH flowers from other species (25-95%, Oakley et al., 2007; Abdala-Roberts et al., 2014; Goodwillie et al., 2018).

Relative investment in seeds of the two flower types can be an important component of overall relative energetic costs of reproduction. In some species such as *Amphicarpaea bracteata* (Schnee and Waller, 1986) and several grasses (Cheplick, 1989; Cheplick, 2023; Jones et al., 2023), there is a seed size heteromorphism with at least some of the CL derived seeds being much larger than CH derived seeds. Additionally, there can be variation in seed number between CH and CL fruits even with similar seed mass (Seguí et al., 2021). However, in *R. humilis* we found that neither seed number per fruit nor average seed weight per fruit differed significantly between CH and CL fruits. This lack of flower type effect on seed weight has been documented in other cleistogamout species such as *Lamium amplexicaule* and *Viola pubescens* (Culley, 2002; Stojanova et al., 2016). Similar seed numbers per fruit between flower types is also not uncommon in other cleistogamous species (Culley, 2002; Albert et al., 2011).

### Total energetic costs of reproduction via the two flower types

We calculated an estimate of the energetic cost of reproduction via CL and CH flowers (expressed as mg/seed) for each population of *Ruellia humilis* using population mean values for the traits described above. Of particular interest is the energetic cost of reproduction via CL flowers relative to CH reproduction (CL/CH). This relative cost of reproduction was as low as around 0.09 in St. Mary’s and Wilmington and up to 0.26 in Sand Ridge. In other words, reproduction via CL flowers represented between 4-10 times less investment in reproductive biomass than reproduction via CH flowers. Only four prior studies, three in a single population, have collected all the necessary data in the same system to calculate relative cost of reproduction of the two flower types (Schemske, 1978; Waller, 1979; Oakley et al., 2007; Winn and Moriuchi, 2009; Seguí et al., 2021), with values ranging from 0.06 to 0.67 (mean = 0.42). The values we estimated for Sand Ridge are comparable (0.26 here vs. 0.38) to those reported for *Impatiens pallida*, (Schemske, 1978). The values for Wilmington and St. Mary’s (around 0.09) however are more similar to the value of 0.06 that we calculated using data from three populations of *Viola jaubertiana* (Seguí et al., 2021). These two populations of *R. humilis* and three populations of *V. jaubertiana* are the clearest examples of the energetic economy of CL flowers to date. However, other studies with large values of relative fertility of CL flowers that would likely yield even lover values of relative cost per seed if all the requisite data for this calculation had been collected.

### Pollen ovule ratio

We measured pollen ovule ratios as another of estimate of energetic costs of the two flower types. Although gametophytes do not contribute much to overall flower mass, there is reason to expect considerable energetic investment in producing pollen and ovules (Schemske, 1978; Schoen and Lloyd, 1984). We found pollen ovule ratios of CL flowers in *R. humilis* were about one third to one half those of CH flowers. Lord (1981) found a similar proportional decrease in P:O of CL flowers of *Lamium amplexicaule*. These findings are consistent with the expectation that CL flowers reduce investment in producing pollen because selfing mechanisms are accurate (Cruden, 1977; Queller, 1984).

### Patterns of relative phenology and degree of chasmogamy

We observed considerable variation among populations in the relative phenology of the two flower types. One thing in common between all three populations is that they do not produce different flower types in distinct seasons as is common in perennial cleistogamous species (Oakley et al., 2007; Winn and Moriuchi, 2009), but see (Seguí et al., 2021; Austin et al., 2022). Instead, Wilmington had a pattern of simultaneous production, whereas Sand Ridge and St. Mary’s had partially overlapping distributions of the two flower types. The lack of distinct seasons for the different flower types suggests that the maintenance of CH flowers is not an adaptive plastic response to producing CH flowers in the season in which they are more favorable (Schoen and Lloyd, 1984; Winn and Moriuchi, 2009). Additionally in all three populations, CH flowers were produced either before or simultaneously with CL flowers, which appears to be uncommon in species with overlapping distributions of the two flower types (Oakley et al., 2007). These populations therefore do not exhibit a “pessimistic strategy” (Cheplick and Quinn, 1982) by investing in CL flower production first, but instead immediately begin investing in the more risky CH flowers. It has also been suggested that CH production is a bet-hedging strategy (Waller, 1980), though this hypothesis is difficult to test. The proportion of total flowers that were chasmogamous (i.e., degree of chasmogamy) was overall around 50% and slightly lower in Sand Ridge than the other two populations.

## Conclusions

Our results provide a clearer picture of the mating system and the selective advantage required to explain the continued maintenance of CH flower production in this species. We estimate that maximum possible population level outcrossing rates are between 43% for Sand Ridge to 61% for St. Mary’s, and considering estimated rates of autogamous selfing in CH flowers brings these maxima down to between 11-33%. An important caveat is that all estimated trait values from our greenhouse common garden experiment may be different if measured under field conditions. Our biomass-based measures of the energetic economy of CL flowers suggest that progeny from CH flowers would have to be 4-10 times more fit to explain their evolutionary maintenance in the face of short-term selection. Reduced relative pollen:ovule ratios in CL compared to CH flowers suggest the selective advantage of CH flowers would need to be even greater than 4-10 fold. Forthcoming work investigates the potential role of both inbreeding depression and heterosis in maintaining CH reproduction (Soto, 2022).

## Acknowledgements

The authors thank Bob Easter from NICHES Land Trust for allowing us to collect seed from Sand Ridge. We also would like to thank Juan Diego Rojas-Gutierrez for assistance with the greenhouse experiment. This work was funded in part by USDA Hatch grant 7000281 to C.G.O via Purdue College of Agriculture, and a Purdue ARGE assistantship to T.Y.S.

---

**Table S1.**
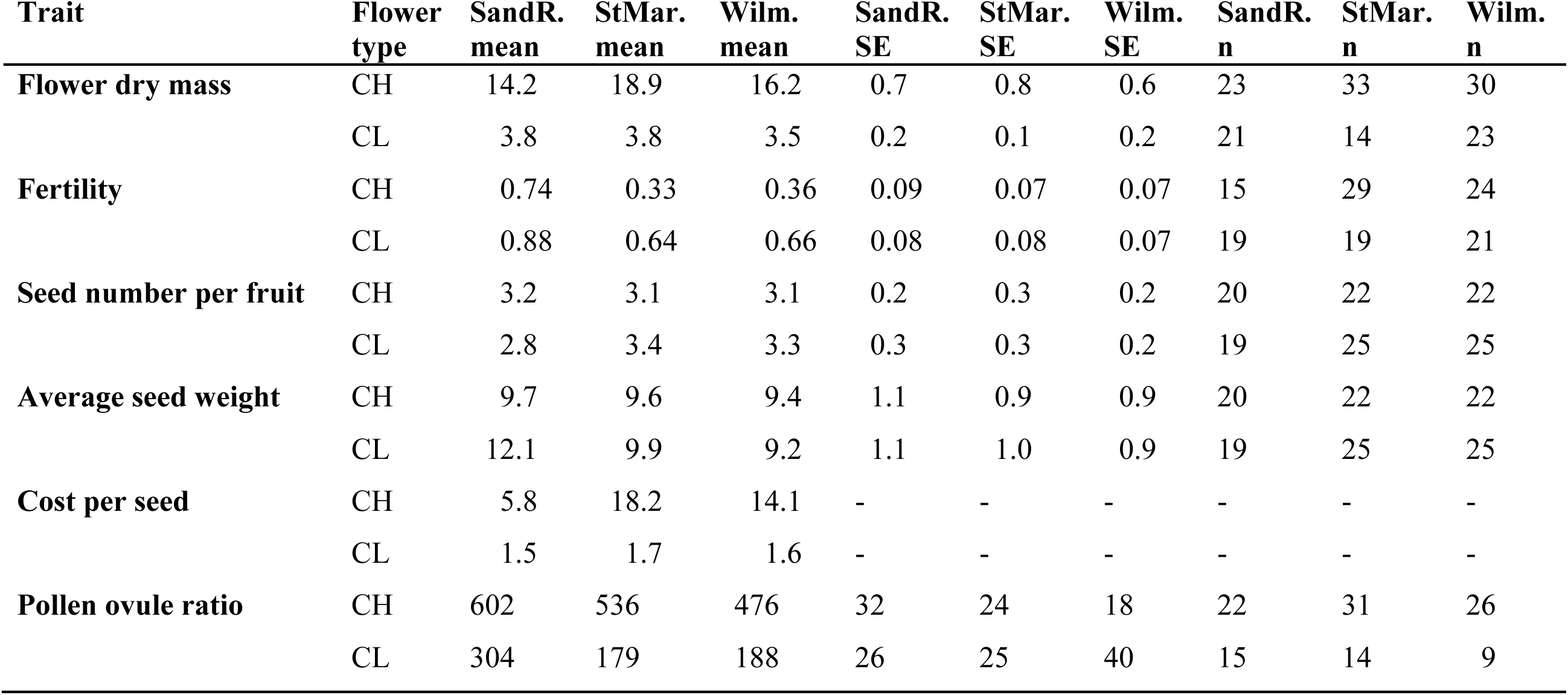
Means, standard errors (SE), and sample sizes (n) for traits for both chasmogamous (CH) and cleistogamous (CL) for the Sand Ridge (SandR.), St. Mary’s (StMar.) and Wilmington (Wilm.) maternal populations. Traits include flower dry mass (mg), fertility, seed number per fruit, average seed weight (μg), cost per seed (mg/seed), and pollen-ovule ratio (P:O). For each population, cost of reproduction of each flower type (mg/seed) was calculated using this formula: [(flower dry mass/fertility) + total seed mass per fruit]/seed number per fruit. Because cost per seed is a population level estimate, no standard error or sample size is given for this trait. For all traits except seed number per fruit, sample size is the number of individual plants sampled. The sample size listed for seed number per fruit and average seed weight represents the number of fruits measured.

## Notes

### Competing Interest Statement

The authors have declared no competing interest.

